# Features fade, pointers persist: dissociable parietal mechanisms in visual working memory formation and maintenance

**DOI:** 10.1101/2025.07.02.662888

**Authors:** Xinchi Yu, Ellen Lau

## Abstract

An emerging body of work has adopted the theoretical construct of pointers: an object is mentally represented by a content-free pointer binding the corresponding features together. Indeed, fMRI studies have highlighted dissociable parietal regions sensitive to pointer and feature load respectively. Specifically, while the superior IPS (intraparietal sulcus) is sensitive to feature load, inferior IPS is only sensitive to the number of pointers – i.e., the number of objects. However, the spatiotemporal dynamics remain unclear, and therefore it is unknown whether these effects reflect visual working memory (VWM) formation, maintenance, or both, especially given limited temporal resolution of fMRI. In our current MEG (magnetoencephalography) study, participants memorized visual arrays with different numbers of objects (object/pointer load: two, four), as well as different features and feature-per-object (color, orientation, bifeatural) in a VWM task. We observed a dissociation between inferior and superior IPS in the temporal dynamics of feature-sensitive and pointer-sensitive responses. While pointer-sensitive signals persisted across VWM formation and maintenance in inferior IPS, feature-sensitivity was only transiently manifested during VWM formation in superior IPS. This spatiotemporal dissociation may reflect a representational architecture optimized for efficiency, reducing the need for sustained neural activity to maintain features once they are bound to pointers. In revealing the spatiotemporal profile of pointer and feature representations, our results provide novel evidence on how pointers underlie energy-efficient neural representations in VWM.

## 1. Introduction

Human visual working memory (VWM), or visual short-term memory, enables us to hold multiple items temporarily, in service of future tasks or computations. This capacity-limited cognitive faculty, remarkably, underlies our extremely flexible cognition and behavior (Ballard et al., 1997; Van Ede & Nobre, 2023; Xie et al., 2020). While it is clear that we can hold multiple (up to three or four) visual objects in VWM (Awh et al., 2007; Cowan, 2001; Zhang & Luck, 2008), the format – or, data structure – of such representation, and how the brain supports this data structure remains a topic of ongoing debate.

One influential proposal is that an object is registered in VWM by a pointer (Awh & Vogel, 2025; Balaban et al., 2019; Ngiam, 2024; Yu & Lau, 2023). These pointers, akin to pointers in programming languages, do not represent features of this object in themselves (Yu, 2025). Rather, a pointer “points” to the representation of corresponding features; the pointers themselves are therefore “content-free” (Jones, Diaz, et al., 2024; Jones, Thyer, et al., 2024; Thyer et al., 2022; Yu & Lau, 2025a). While this proposal was originally conceived as an algorithmic-level theory (Kahneman et al., 1992; Pylyshyn, 2001), recent decades have seen significant advances towards understanding how the brain implements these “pointers”. A groundbreaking fMRI study on VWM showed that inferior IPS (intraparietal sulcus) hosts the representation of pointers with univariate neural responses (Xu & Chun, 2006): univariate BOLD signal in this region increases with the number of objects, plateauing beyond the capacity limit of three to four items, and is insensitive to the number of features per object to be held. On the other hand, superior IPS is sensitive to the number of features – not (just) the number of objects.Qualitatively similar findings with fMRI have been obtained in other studies (Bettencourt & Xu, 2016; Naughtin et al., 2016; Xu, 2009), yet several questions remain.

First, due to the sluggish temporal resolution of fMRI, it remains unclear whether these reported effects only reflect perception or VWM formation processes (cf. Xie & Zhang, 2018), or whether such effects extend to VWM maintenance. Second, one prediction following the pointer hypothesis is that, as sustaining neural activations is energy-consuming, the best engineering solution is to only sustain the representation for the pointers but not necessarily the features to which they “point”. The addressing of these features may rather be accomplished by means of activity-silent neural mechanisms (e.g., via synaptic or intracellular encoding; Gallistel & King, 2009; Rose et al., 2016; Wolff et al., 2015). This hypothesis is in line with anecdotal reports that, in contrast to the case for VWM formation, reliable neural decoding of features during VWM maintenance is surprisingly weak or even absent (Bae & Luck, 2018; Teichmann et al., 2022; Yeh et al., 2025). However, direct evidence for this potential temporal dissociation between pointers and features remains lacking. To sum up, prior fMRI studies on VWM representation of pointers and features leave open at least three possible spatiotemporal dynamics (**Figure 1A**):

**Figure 1.**
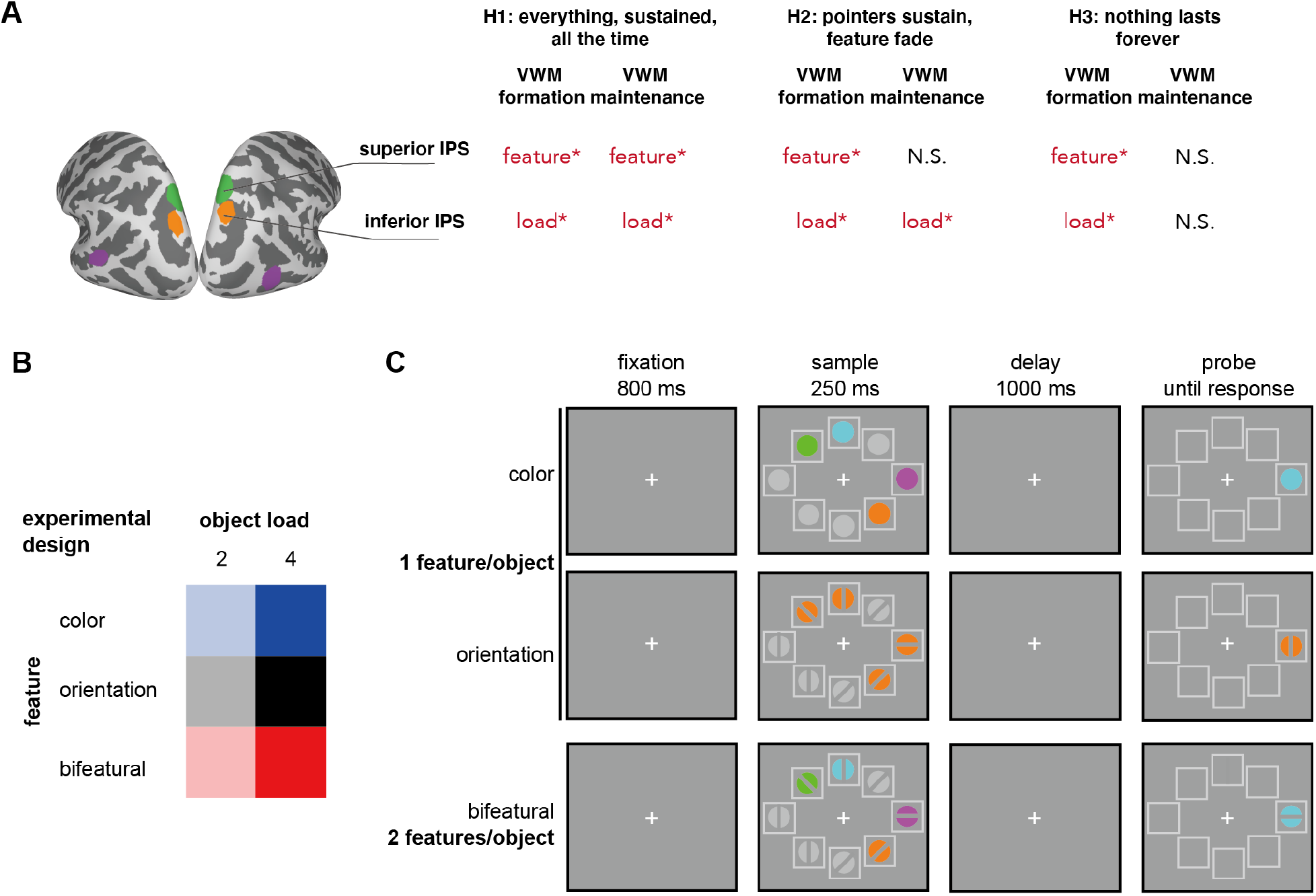
**(A)** Three literature-informed hypotheses on the spatiotemporal dynamics across inferior and superior IPS. **(B)** Experimental conditions: we employed a 2 × 3 design, with two object load conditions (two vs. four objects), and three feature conditions (color-only, orientation-only, and bifeatural). **(C)** Illustration of a trial across three feature conditions with the example of object load = 4.

(H1) **Everything, sustained, all the time**: inferior IPS holds pointer-sensitive signal across VWM formation and maintenance, while superior IPS is sensitive to features across time as well.

(H2) **Pointers sustain, features fade**: while inferior IPS holds the representation of pointers across VWM formation and maintenance, sensitivity of superior IPS to features is weaker beyond VWM formation.

(H3) **Nothing lasts forever**: prior fMRI findings of the labor division between inferior and superior IPS only reflected VWM formation but not maintenance, so that both pointer-sensitive and feature-sensitive effects were only constrained to VWM formation.

To evaluate these hypotheses, in the current study we sought to uncover the temporal dynamics in inferior and superior parietal regions, enabled by the combination of good spatial and temporal resolutions in MEG (magnetoencephalography). Complementarily, we also examined the temporal dynamics of LOC (lateral occipital cortex), which was observed to be sensitive to feature load in some of the aforementioned fMRI studies albeit was not reported in all cases (Xu & Chun, 2006).

We employed a 2 × 3 design, with two object load (“load” in short) conditions (two vs. four objects), and three feature conditions (color-only, orientation-only, and bifeatural [both color and orientation]; **Figure 1B**). In color trials and orientation trials, only color or orientation needs to be encoded for each object, while in bifeatural trials, both color and orientation need to be encoded for each object.

## 2. Materials and Methods

Data and processing scripts for this study will be openly shared on OSF upon acceptance.

### 2.1 Participants

27 participants (age 19-26, 14 female) entered our data analysis. One additional participant was excluded because of chance level (49%) performance in some blocks. The participants received monetary reimbursement or course credit for their participation. The procedures are approved by the Institutional Review Board of University of Maryland College Park.

### 2.2 Procedure

Each participant was presented with 600 trials: 200 color trials, 200 orientation trials, and 200 bifeatural trials. Within each feature condition (color, orientation, bifeatural), half of the trials had a load of two objects and half had a load of four objects. These 600 trials were evenly divided into six blocks: two color blocks, two orientation blocks, and two bifeatural blocks. Each block contained 100 trials, and the order of the blocks was randomized for each participant, with the constraint that each type of block appeared once in the first three and last three blocks respectively. In other words, the trials in each block belonged to the same feature condition (color, orientation, bifeatural) but were intermixed in terms of memory load (two vs. four objects).

In each trial, the participants were first presented with a fixation cross with a duration of 800 ms against a grey background, followed by a sample memory array with a duration of 250 ms (see **Figure 1C**). The sample consisted of two or four unique target objects. For the color trials, the colors were sampled without replacement from the following seven colors: red [255,0,0], green [0,255,0], blue [0,0,255], yellow [255,255,0], purple [255,0,255], teal [0,255,255], and orange [255,128,0]. For the orientation trials, the bar orientation was sampled without replacement from the following four orientations: 0°, 45°, 90°, 135°. The color of the disk behind these target orientations was randomly sampled from the above colors for each participant and was kept consistent throughout that experimental session for that participant. For bifeatural trials, each target object was a combination of a color and an orientation sampled in the aforementioned way. The target objects (2.1° × 2.1°) occupied two or four out of all eight boxes (light gray [191,191,191], 2.5° × 2.5° each), located at eight fixed locations evenly distributed 4.5° from the center of the screen. The box contours were employed to discourage grouping (Xie & Zhang, 2017). Filler gray disks ([166,166,166]) occupied the rest of the locations for all trials, additionally with an orientation bar at a random orientation for the orientation and bifeatural trials. The orientation bars had a width of 0.5°. A blank screen delay of 1000 ms followed the sample, after which the probe appeared on the screen.

For the probe, only one object was presented at one of the locations previously occupied by a target object. Half of the trials had a match probe, and the other half had a change probe. On “match” trials, this object was the same one as the one in the sample for that location; on “change” trials, this probe object was randomly selected from other target features used in this block. For example, for a change trial in the color condition, participants were probed with a color that appeared in the sample but not at the probed location. For a change trial in the bifeatural condition, participants were probed with an object with the right color for that location but a wrong – yet previously presented – orientation (or the right orientation yet a wrong color). This set up required the participants to represent bindings across features instead of free-floating features (Yu & Lau, 2025b), as feature binding is hypothesized to be one of the key functions of pointers (Yu & Lau, 2023). Participants pressed one button to indicate “match”, and pressed another button to indicate “change” (buttons randomized for each participant), with no time limit for response. After their response, they were shown a line of asterisks on the screen, where they would press any button to proceed to the next trial in a self-paced manner. Prior to the experiment, participants were instructed to blink only during this period, and to avoid blinking during the trials. While the fixation cross was presented on the screen, the participants were instructed to fixate at it. They were also instructed against verbal rehearsal. At least 24 practice trials were run for each participant prior to the experiment. The setup was broadly consistent with the EEG study of Thyer et al. (2022), and the experiment was run in MATLAB R2009a with Psychtoolbox 3 (Brainard, 1997; Kleiner et al., 2007).

### 2.3 Behavioral analyses

Accuracy was calculated for each participant for each condition. The responses were subject to a 2 × 3 repeated-measures ANOVA (load: two, four × feature: color, orientation, bifeatural; **Figure 1B**). Significant interactions were followed up with pairwise comparisons using t-tests The statistics reported in the current article are all based on two-tailed tests.

### 2.4 MEG recording

Prior to recording, five head position indicator coils were affixed to each participant’s head, and the position of these coils relative to the nasion and tragus, as well as the participant’s head-shape, was digitized using a Polhemus 3SPACE FASTRAK system in order to determine the participant’s accurate placement in the MEG dewar. During the experimental sessions, participants laid supine in a dimly lit magnetically shielded room (Vacuumschmelze GmbH & Co. KG, Hanau, Germany). Continuous MEG recording was executed using a 160-channel axial gradiometer whole-head system (Kanazawa Institute of Technology, Kanazawa, Japan), and data was sampled at 1000 Hz (60 Hz online notch filter, DC-200 Hz recording bandwidth).

### 2.5 MEG analyses

#### 2.5.1 Sensor-level analyses

MEG sensor-level data was analyzed by customized code in MATLAB R2020b with MNE-Python 1.5.1 (Gramfort et al., 2013), with a similar procedure as (Yu & Lau, 2025a). First, noisy and flat channels were identified for each participant with Maxwell filtering, and these channels (1-3 per participant) were interpolated with the MNE algorithm. Then the environmental magnetic interferences were suppressed using temporal signal space separation (tSSS; Taulu & Simola, 2006). A low-pass infinite impulse response (IIR) filter with an upper cutoff frequency of 40 Hz was then applied to the MEG data. Ocular and cardiac artifacts were removed using independent component analysis (ICA; fastica algorithm); 2-7 ICA components were removed for each participant. Epochs of −200:1200ms time-locked to the presentation of sample stimuli in each trial were extracted. The 200 ms pre-stimulus interval was used as the baseline interval. Trials with a maximum peak-to-peak signal amplitude higher than 3000 fT were excluded. For each participant, the number of epochs per condition was matched with the constraint of minimizing timing differences, using the mintime algorithm. This resulted in 88-100 epochs/condition entering further data analysis for each participant (that is, 88-100% of all trials, mean = 98.81%). For each participant and each condition (i.e., a certain load condition in a certain feature block), we calculated the mean event-related field (ERF) response for each channel. Following Yu & Lau (2025), the RPA (right posterior activity) ROI was defined with the following channels: MEG054, MEG055, MEG071. Mean ERFs were calculated within this ROI. Comparisons across conditions were statistically evaluated with repeated-measures ANOVA in JASP 0.18.1 (JASP Team, 2023).

#### 2.5.2 Source space analyses

For the source space analyses, the “fsaverage” template brain from the FreeSurfer software package (http://surfer.nmr.mgh.harvard.edu) was used as an approximation for individual anatomy (four-fold icosahedral subdivision), and digitizer data was used to manually coregister individual head position to the template brain. A three-compartment boundary element model computed for this template brain with the linear collocation approach was used in the forward calculation (Hämäläinen & Sarvas, 1989; Mosher et al., 1999). Then we created an inverse operator, for the averaged responses for each participant-condition combination. Noise covariance estimates were derived from data recorded in the −200:0 ms baseline period. The orientation of the dipoles was subjected to a loose constraint with weighting factor 0.2 (Lin et al., 2006), and depth was weighted at 0.8. Then the inverse operator was applied to the corresponding averaged evoked responses for the corresponding participant and condition, with regularization λ = 1/9 and the dSPM algorithm (Dale et al., 2000). Individual estimates were then smoothed using seven iterative steps to spread estimated activity to neighboring vertices.

The source space ROIs were defined a priori based on the fMRI study of Xu & Chun (2006), converting their original Talairach coordinates to MNI coordinates (Papademetris et al., 2006): inferior IPS, left [−25,−74,30], right [25,−69,36]; superior IPS, left [−14,−73,51], right [15,−72,55]; LOC, left [−46,−50,4], right [41,−68,2]. Each ROI was grown from the nearest vertex to these MNI coordinates, within a maximum radius of 10 mm on the white matter surface based on geodesic distance, constrained to the corresponding hemisphere. The responses in each ROI were averaged across two consecutive time windows respectively: 200-400 ms (VWM formation) and 400-1200 ms (VWM maintenance). For each time window, the responses were subject to a 2 × 3 repeated-measures ANOVA (load: two, four × feature: color, orientation, bifeatural). For visual examination, a contrast map based on t-values was created by comparing across load conditions (paired t-test) vertex by vertex with a threshold of α = 0.05 across a series of 200 ms time windows from 200 ms to 1200 ms (**Figure S1**).

## 3. Results

### 3.1 Behavioral results

For each of the six conditions, participants had above-chance performance, with bootstrapped 95% CI as follows: color, 2 (93.1%-96.0%), color, 4 (77.2%-85.1%), orientation, 2(83.9%-89.7%), orientation, 4 (66.4%-71.7%), bifeatural, 2 (84.6%-89.1%), bifeatural, 4 (64.9%-69.6%). A 2 (load: two, four) × 3 (feature: color, orientation, bifeatural) repeated-measures ANOVA revealed a significant main effect of load, *F*(1,26) = 403.28, *p* < 0.001, *η*_*p*_^2^ = 0.94, a significant main effect of feature, *F*(2,52) = 81.96, *p* < 0.001, *η*_*p*_^2^ = 0.76, as well as a significant interaction effect, *F*(2,52) = 5.37, *p* = 0.008, *η*_*p*_^2^ = 0.17. The interaction effect appeared to be driven by a larger numerical impact of load on orientation and bifeatural trials compared to the color trials: comparing the drop in accuracy across load 2 and 4 conditions, we observed a significant difference between color and orientation trials (paired t test, *t*(26) = −2.13, *p* = 0.04, Cohen’s d = −0.41), and a significant difference between color and bifeatural trials (*t*(26) = −2.98, *p* = 0.006, Cohen’s d = −0.57), but not a significant difference between orientation and bifeatural trials (*t*(26) = −1.07, *p* = 0.30, Cohen’s d = −0.21). While the magnitude of the load effect was thus greatest in orientation and bifeatural trials, color conditions still showed a robust, statistically significant load effect in their own right (*t*(26) = 7.89, *p* < 0.001, Cohen’s d = 1.52).

### 3.2 MEG sensor-level results

Similar to our prior study (Yu & Lau, 2025a), we analyzed the ERF responses in the 400-1200 ms time window in right posterior sensors (the RPA ROI). A 2 (load: two, four) × 3(feature: color, orientation, bifeatural) repeated-measures ANOVA revealed a significant main effect of load, *F*(1,26) = 21.88, *p* < 0.001, *η*_*p*_^2^ = 0.46, as well as a significant interaction effect, *F*(2,52) = 4.39, *p* = 0.02, *η*_*p*_^2^ = 0.14. The main effect of feature was not statistically significant, *F*(2,52) = 1.51, *p* = 0.23, *η*_*p*_^2^ = 0.06. See Figure 2.

**Figure 2.**
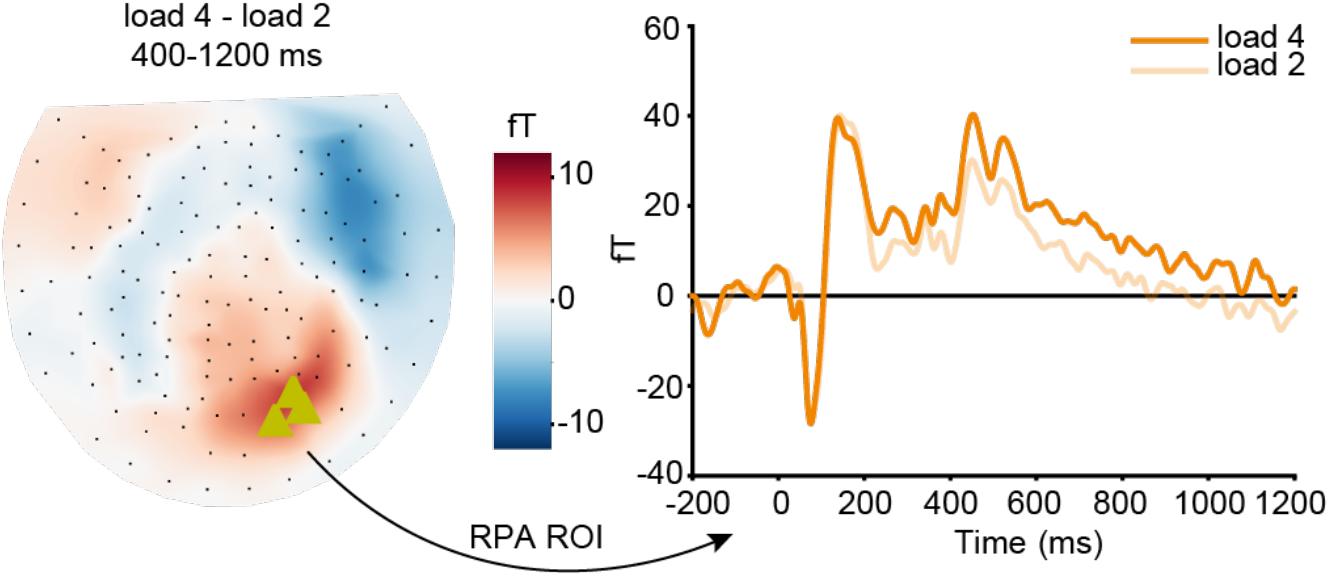
ERF (event-related field) responses at the RPA ROI across the two load conditions. RPA: right posterior activity; ROI: region-of-interest.

Follow-up paired t-tests across two load conditions revealed a significant difference for color trials, *t*(26) = −4.23, *p* < 0.001, Cohen’s d = −0.82, as well as for the bifeatural trials, *t*(26) =−3.19, *p* = 0.004, Cohen’s d = −0.61. The difference was only numerical for the orientation trials, *t*(26) = −1.46, *p* = 0.16, Cohen’s d = −0.28. These results are largely in line with prior reports that RPA is sensitive to the number of pointers but not the number of features (Yu & Lau, 2025a).

### 3.3. MEG source-level results

For inferior IPS, a main effect of load emerged for VWM formation (*F*(1,26) = 4.13, *p* = 0.05, *η*_*p*_^2^ = 0.14), and extended into VWM maintenance (*F*(1,26) = 5.92, *p* = 0.02, *η*_*p*_^2^ = 0.19). In contrast, no effects of feature were observed in this region. For VWM formation, neither the main effect of feature nor the interaction effect was statistically significant: feature, *F*(2,52) = 0.60, *p* = 0.55, *η*_*p*_^2^ = 0.02; interaction effect, *F*(2,52) = 0.80, *p* = 0.46, *η*_*p*_^2^ = 0.03. During VWM maintenance, neither the main effect of feature nor the interaction effect was statistically significant: feature, *F*(2,52) = 0.66, *p* = 0.52, *η*_*p*_^2^ = 0.03; interaction effect, *F*(2,52) = 0.40, *p* = 0.67, *η*_*p*_^2^ = 0.02. See Figure 3.

**Figure 3.**
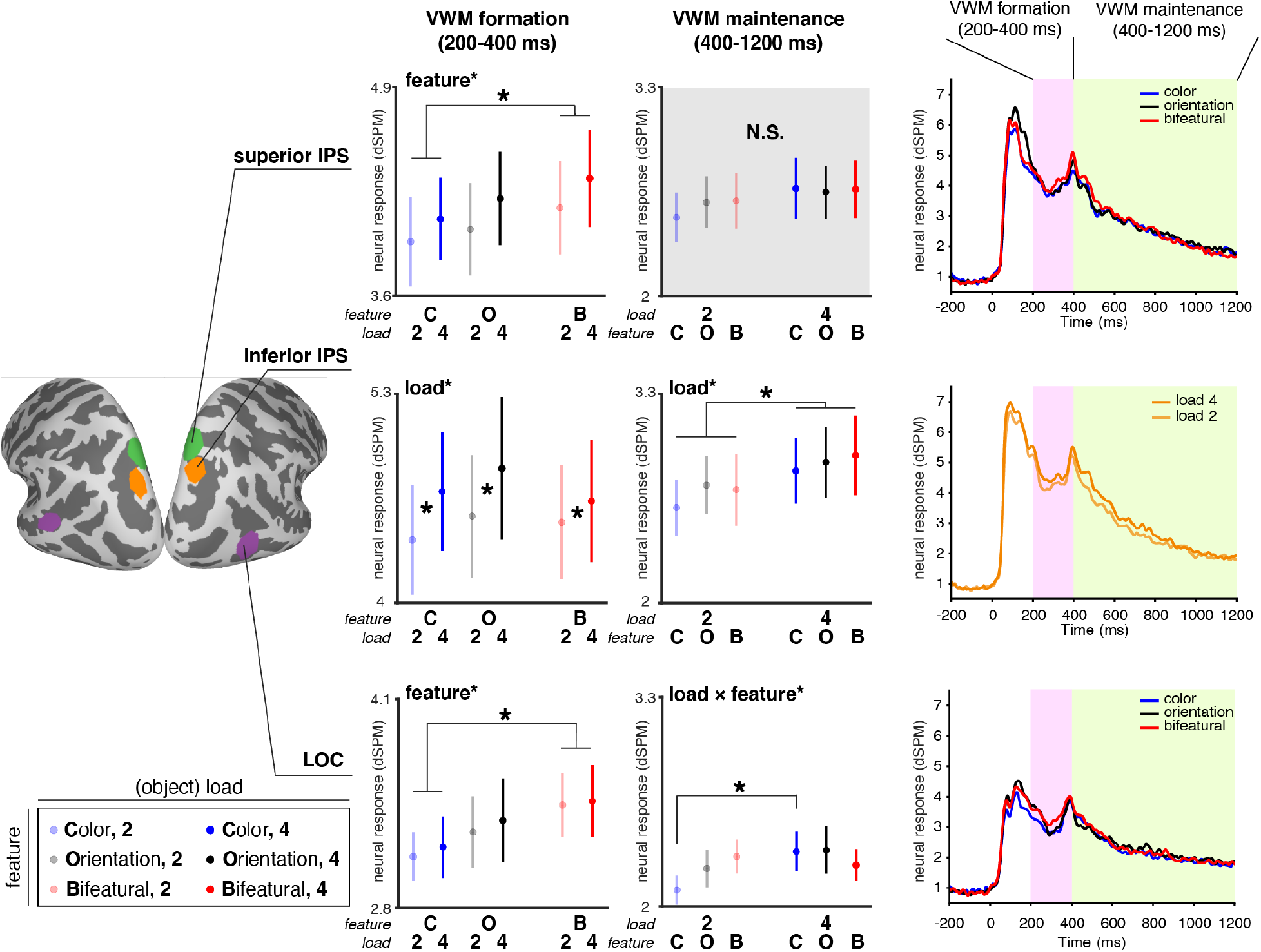
Leftmost and middle columns: responses during VWM formation (200-400 ms) and maintenance (400-1200 ms) in three a priori ROIs: inferior IPS, superior IPS and LOC. The rightmost column demonstrates neural responses (dSPM) across time in the corresponding ROI grouped by feature or object load conditions. Error bars stand for standard error. *: *p* ≤ 0.05.VWM: visual working memory; ROI: region-of-interest; IPS: intraparietal sulcus; LOC: lateral occipital cortex; C: color; O: orientation; B: bifeatural; 2: object load = 2; 4: object load = 4.

For superior IPS, a main effect of feature manifested for VWM formation (*F*(2,52) = 3.60, *p* = 0.03, *η*_*p*_^2^ = 0.12. The main effect of load was only marginally significant, *F*(1,26) = 3.61, *p* = 0.07, *η*_*p*_^2^ = 0.12, and the interaction effect was non-significant, *F*(2,52) = 0.09, *p* = 0.91, *η*_*p*_^2^ = 0.004. Post hoc pairwise comparison (Bonferroni-corrected) across the three feature conditions revealed a significant difference between color and bifeatural conditions, *t*(26) = −2.68, *p* = 0.03, Cohen’s d = −0.16. The differences between the other two pairs of feature conditions were not statistically significant. During VWM maintenance, none of the effects were statistically significant (F values < 2.48, p values > 0.12).

For LOC, a main effect of feature manifested for VWM formation (*F*(2,52) = 3.44, *p* = 0.04, *η*_*p*_^2^ = 0.12. Neither the main effect of load nor the interaction effect was significant: load, *F*(1,26) = 0.48, *p* = 0.50, *η*_*p*_^2^ = 0.02; the interaction effect, *F*(2,52) = 0.11, *p* = 0.89, *η*_*p*_^2^ = 0.004. Post hoc pairwise comparison (Bonferroni-corrected) across the three feature conditions revealed a significant difference between color and bifeatural conditions, *t*(26) = −2.62, *p* = 0.03, Cohen’s d = −0.28. The differences between the other two pairs of feature conditions were not statistically significant. During VWM maintenance, there was a significant interaction effect between feature and load, *F*(2,52) = 3.18, *p* = 0.05, *η*_*p*_^2^ = 0.11. The main effect of load was marginally significant, *F*(1,26) = 3.48, *p* = 0.07, *η*_*p*_^2^ = 0.12, and the main effect of feature was not significant, *F*(2,52) = 0.37, *p* = 0.69, *η*_*p*_^2^ = 0.01. The interaction effect was driven by differential impact of load on different feature conditions: for color trials, paired t-test (Bonferroni-corrected) revealed a significant difference across the two load conditions, *t*(26) = −2.63, *p* = 0.04, Cohen’s d = −0.38. The load-driven difference was not statistically significant for orientation trials or bifeatural trials (|t values| < 2, Bonferroni-corrected p values > 0.18).

## 4. Discussion

In the current study, we observed a spatiotemporal dissociation of the representation of pointers and features in VWM. Namely, pointer-sensitive representations were discernable across both VWM formation (200-400 ms) and maintenance (400-1200 ms) in inferior IPS, while feature-sensitive signals were transiently observable during VWM formation but not maintenance in superior IPS. Taken together, our Hypothesis 2 was supported: that pointers persist through sustained neural activities across VWM formation and maintenance, while activity-silent mechanisms are sufficient for recovering the features to which they point.

Complementarily, feature-sensitive signals were also detected in LOC during VWM formation. However, there appeared to be some change in representational geometry across experimental conditions when progressing into VWM maintenance: the representation during VWM formation was dominated by the number of features per object, while the representation during VWM maintenance showed a significant interaction effect between feature and load, as well as a marginally-significant main effect of load. Follow-up pairwise comparison revealed that the effects appeared to be primarily driven by a difference within color trials, across load two and four conditions.

### 4.1 Functional dissociation across parietal patches for VWM

Although the involvement of posterior parietal cortex in visual working memory has been well-established (Becke et al., 2015; Chopurian et al., 2024; Lorenc et al., 2018; Todd & Marois, 2004; Xu & Chun, 2006), the division of labor within this region has not been fully charted (Brigadoi et al., 2017). Extending prior fMRI reports suggesting a spatial dissociation where inferior IPS represents pointers and superior IPS represents features (Xu, 2009; Xu & Chun, 2006), our current results support a *spatiotemporal* dissociation across these two posterior parietal patches.

Such results are in line with the function of pointers: in pointing to the features for a specific object, they provide the infrastructure for feature binding (Awh & Vogel, 2025; Yu & Lau, 2023). If that were the case, sustained firing in the neural population coding the features themselves would not be necessary during VWM maintenance, as long as the representation of pointers is sustained, given that the pointer “has the addresses” of the corresponding features (Green & Quilty-Dunn, 2021). Representing pointer-feature correspondence is feasible via mechanisms other than neural firing, for example, via so-called activity-silent mechanisms. Two major possibilities have been proposed in the literature as the neural substrate of activity-silent mechanisms: one argues for a pivotal role of synaptic connectivities (Rose et al., 2016; Wolff et al., 2015), and the other theory highlights the capacity of memory storage within neurons through molecular mechanisms (Akhlaghpour, 2022; Gallistel & King, 2009). Future studies should aim at elucidating the format of activity-silent representations in VWM.

### 4.2. Temporal progression of representations in the LOC

Another curious finding in the current study was the temporal progression of the representational geometry across the two consecutive time windows in LOC. During VWM formation, the responses were dominated by a main effect of feature: regardless of object load, the bifeatural trials elicited higher responses compared to color-only trials. This representational geometry then, during VWM maintenance, shifted to a marginally significant effect of object load and a significant interaction effect between feature and object load. As inferior IPS demonstrates clear load-dependent responses across VWM formation and maintenance, this result is in line with recent fMRI reports that representations in lateral occipital regions is progressively shaped by the representational geometry in parietal regions across time (Xu, 2023). In the fMRI study of Xu (2023), participants memorized real-world objects in a visual working memory task, where in each trial the objects belong to a certain semantic category (e.g., houses, cars). It was observed that the representational geometry across object categories in lateral occipital regions became more aligned with that in inferior parietal regions during VWM maintenance compared to during formation. Our current results extended the observation of Xu (2023) from real-world objects to relatively meaningless colors and orientations, suggesting a potentially omnipresent parietal-LOC dynamic relationship regardless of whether long-term semantic knowledge is consulted. According to this hypothesis, the representational geometry in LOC is dynamically warped by that in parietal regions across VWM formation and maintenance. While an emerging body of research highlights representational and processing differences across less meaningful vs. more meaningful objects (Chung et al., 2024; Li et al., 2025; Yu et al., 2025), our observations suggest that this parietal-LOC dynamics may be shared across visual working memory tasks regardless of object meaningfulness.

## 5. Conclusion

In sum, these results support a critical role for inferior IPS in instantiating pointers across VWM formation and maintenance, compensating for a less “active” representation for features in superior IPS and LOC during VWM maintenance. These results also indicate that MEG, even with its lower spatial resolution than fMRI, is able to resolve activity in the parietal cortex from subtle neighboring regions with different functional roles. These results thus offer new neural markers – for example inferior IPS activities tracking the number of pointers – for investigating visual working memory with improved spatiotemporal precision.

## Author contributions

Xinchi Yu: conceptualization, methodology, software, formal analysis, investigation, data curation, writing - original draft, writing - review & editing, visualization

Ellen Lau: conceptualization, methodology, resources, writing - review & editing, supervision, funding acquisition

## Acknowledgements

We would like to thank Audrey Laun for technical support. The study was supported by funding from the Department of Linguistics, University of Maryland.

