## Supplementary material for "Features fade, pointers persist: dissociable parietal mechanisms in visual working memory formation and maintenance": Figure S1

### Supplementary Materials

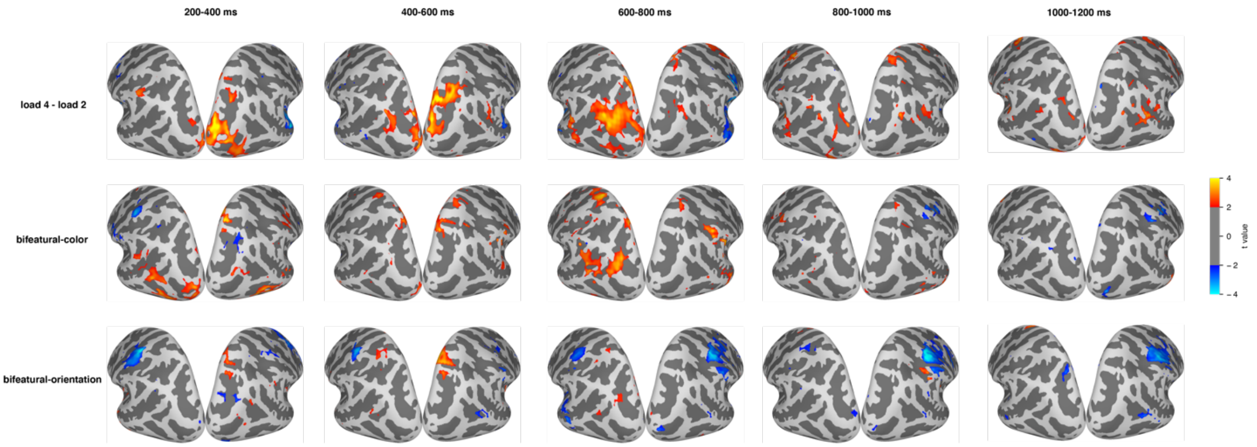


**Figure S1.** Visualization of activation differences across object load conditions 4 vs. 2, as well as each pair of feature conditions, in 200 ms time bins from 200 ms to 1200 ms. Only vertices that reached $p < 0.05$ (paired t-test, two-tailed) are shown.
